# Swine Promyelocytic Leukemia Isoform II Inhibits Pseudorabies Virus Infection by Suppressing Viral Gene Transcription in PML-NBs

**DOI:** 10.1101/2020.03.23.004697

**Authors:** Cuilian Yu, Aotian Xu, Yue Lang, Chao Qin, Xiufang Yuan, Wenhai Feng, Mengdong Wang, Chao Gao, Jinwen Chen, Rui Zhang, Jun Tang

## Abstract

Promyelocytic leukaemia nuclear bodies (PML-NBs) possess an important intrinsic antiviral activity against α-herpesvirus infection. PML is the structural backbone of NBs, comprising different isoforms. However, the contribution of each isoform to α-herpesvirus restriction is not well understood. Here, we report the role of PML-NBs and swine PML (sPML) isoforms in pseudorabies virus (PRV) infection in its natural host swine cells. We found that sPML-NBs exhibit an anti-PRV activity in the context of increasing the expression level of endogenous sPML. Of four sPML isoforms cloned and examined, only isoform sPML-II/IIa, not sPML-I and IVa, expressed in a sPML knockout cells inhibits PRV infection. Both the unique 7b region of sPML-II and sumoylation-dependent normal formation of PML-NBs are required. 7b possesses a transcriptional repression activity and suppresses viral gene transcription during PRV infection with the cysteine residue 589 and 599 being critically involved. We conclude that sPML-NBs inhibit PRV infection by repressing viral gene transcription through the 7b region of sPML-II.

**IMPORTANCE:** PML-NBs are nuclear sites that mediate the antiviral restriction of α-herpesvirus gene expression and replication. However, the contrition of each PML isoform to this activity of PML-NBs is not well characterized. Using PRV and its natural host swine cells as a system, we have discovered that the unique C-terminus of sPML isoform II is required for PML-NBs to inhibit PRV infection by directly engaging in repression of viral gene transcription. Our study not only confirms in swine cells that PML-NBs have an anti-viral function, but also presents a mechanism to suggest that PML-NBs inhibit viral infection in an isoform specific manner.

## INTRODUCTION

Promyelocytic leukaemia nuclear bodies (PML-NBs) is a PML protein-based sub-nuclear structure with an intrinsic anti-viral activity against a wide range of RNA and DNA viruses. More than 150 proteins have been identified either transiently or residentially associated with PML-NBs, many of which possesses an anti-viral activity including Sp100, Daxx, ATRX, HIRA and MORC3 (1-7). As a countermeasure, numerous viruses have evolved strategies to disrupt or degrade PML-NBs underscoring the importance of this structure in cellular anti-viral defense. In the case of α-herpesvirus HSV-1, PML is ubiquitinated and degraded by the viral E3 ubiquitin ligase ICP0 (8). Thus, an ICP0-deleted HSV-1 mutant is often used to study the role of PML-NBs in HSV-1 restriction.

PML-NBs are involved in multiple mechanisms to restrict herpesvirus infection. It has been long known that PML-NBs have a complex and intimate relationship with herpesvirus DNAs. Recent studies have shown that in addition to immediately entrapping a HSV-1 viral genome after nuclear entry and blocking its replication, PML-NBs are also recruited to the sites of progeny viral DNAs by the nuclear DNA sensor IFI16, and contribute to the repression of viral gene transcription (9-11). The involvement of PML-NBs in this process concerns PML-NBs associated proteins. For example, Daxx and ATRX are critically involved in an epigenetically silencing mechanism (1, 5, 12, 13). However, it is not clear whether PML protein plays a direct role in transcription repression in addition to the recruitment of other proteins. In addition, PML-NBs are also engaged in eliciting innate immune responsive gene transcription, as well as the sequestration of a viral capsid protein to restrict virus infection (14-17).

PML is the structural component of PML-NBs, which belongs to the tripartite motif family with a characteristic RBCC region that includes a RING domain, two B boxes and a coiled-coil (CC) region (18, 19). Due to alternative splicing of mRNAs, a single *PML* gene generates six major nuclear isoforms, referred as PML-I to PML-VI in human cells. These isoforms share a common N-terminal region (exon 1 to 6) containing the RBCC motif (exon 2 to 3), but differ in their C-termini (18, 20, 21). Derivatives of each isoform lacking exon 5 (referred as a) or exon 5 and 6 (referred as b) or exon 4, 5 and 6 (referred as c) might also exist (18, 20). Increasing evidence indicate the unique C-terminal region of each isoform contributes greatly to the composition and functionality of PML-NBs (15, 20, 22, 23), however, the exact role of each isoform remains poorly understood. In the context of anti-herpesvirus defense, it has been reported that in varicella-zoster virus (VZV) infection only PML-IV sequesters the capsid protein encoded by ORF23 leading to VZV restriction (16), whereas in HSV-1 infection PML-I and PML-II play a major role with an unknown mechanism (24).

Herpesvirus infection is considered to be species specific in general, which is partly as a result of long-term co-evolution between viruses and host species. However, the relationship between herpesvirus infection and PML-NBs are mostly characterized in human cells. Given the complexity of PML isoforms and their differential role in VZV and HSV-1 infection in human cells, we hypothesize that PML-NBs in other species might also evolve a specialized function targeting their natural herpesviruses in an isoform specific manner. In this study, we have explored PML-NBs in swine cells in relation to pseudorabies virus (PRV) infection, particularly the role of swine PML (sPML) isoform in restricting PRV infection, and found that sPML-NBs inhibits PRV infection in a sPML-II dependent manner.

PRV is a swine α-herpesvirus which can cause Aujeszky’s disease characterized by respiratory distress, nervous disorder and reproductive failure (25). It is often used as a complement model to study the life-cycle and pathogenicity of alpha-herpesvirus subfamily. As with HSV-1, PRV infection in human cells results in disappearance of PML-NBs. EP0, the PRV ortholog of ICP0, degrades PML in human cells (26, 27). However, the relationship between PRV infection and PML-NBs in its natural host swine cells has not been characterized. In this study, we found that PML-NBs in swine cells were disrupted during PRV infection and that the anti-PRV activity of sPML-NBs may depend on the expression level of sPML. Of four sPML isoforms cloned and examined, only isoform II/IIa which contain exon 7b restrict PRV infection. Exon 7b has an ability to repress viral gene transcriptions with cysteine residues in a ring-like region being critically involved.

## MATERIALS AND METHODS

### Cell culture and viruses

HEK293T cells (human embryonic kidney, ATCC #CRL-3216), PK15 cells (porcine kidney cells, ATCC #CCL-33), Vero cells (ATCC #CCL-81) and CRL cells (porcine alveolar macrophage cells) were cultured in Dulbecco’s modified Eagle’s medium (DMEM). Primary porcine alveolar macrophages (PAM) cells were obtained by lavaging the lungs of 6–8-wk-old specific pathogen–free (SPF) pigs, as described previously (28), and maintained in RPMI 1640. The primary porcine kidney cells were harvested from the kidneys of 21 day-old specific pathogen–free (SPF) piglets, as described previously (29), and maintained in DMEM. All cells were cultured in medium supplemented with 10% (v/v) FBS and maintained in a humidified incubator with 5% CO2 at 37°C.

PRV WT (Bartha K61), the recombinant PRV EP0-Knockout virus (PRV-EP0 KO) and KOS strain of HSV-1 were described previously (30, 31).

### Reagents

Anti-GFP (SC-9996) antibody was purchased from Santa Cruz Biotechnology (Santa Cruz, CA, USA). FLAG (M2 F-1804) antibodies, Triton X-100 and N-ethylmaleimide (NEM) were purchased from Sigma (St Louis, MO, USA). Anti-α-Tubulin mAb PM054 was purchased from MBL. DAPI (4,6-diamidino-2-phenylindole) was purchased from Beyotime Institute of Biotechnology. The antibodies against PRV TK, PRV US3, PRV EP0 were described previously (30, 32, 33). Mouse polyclonal antibodies against PRV IE180, VP5 and gD were raised in mice individually with the 1-666aa, 710-1280aa, 240-400aa region of each protein as antigens. Rabbit polyclonal antibodies against swine PML were described previously (33). Sodium dodecyl sulfate (SDS) was purchased from Scientific Research Ievei. DL-Dithiothreitol (DTT) and bovine serum albumin fraction (BSA) were purchased from Amresco Biotechnology. Puromycin and polybrene were also purchased from Amresco. Swine IFNα (sIFNα) was described previously (33).

### Plasmids and transfection

sPML-I, -II, -IIa and -IVa cDNAs were amplified by PCR using cDNAs made from sIFNα-stimulated PK15 cells as templates and then cloned into Flag-/pRK5, Flag-/pSin-EF2-puro and Flag-GFP-/pSin-EF2-puro vector. All the mutants including deletions, point mutations and fusions were generated by PCR and cloned into Flag-/pRK5 or pSin-EF2-puro vector or both. 7b-nls was created by fusing the NLS sequence of 5’-PKKKRKV-3’ to the C-terminus of 7b. Gal-/pRK5, 5xGal-TK-luciferase reporter and pCMV-β-galactosidase plasmid were previously described (34). All the constructs were confirmed by DNA sequencing.

List of PCR and mutagenic primer is provided in Table 1.

**TABLE 1:**
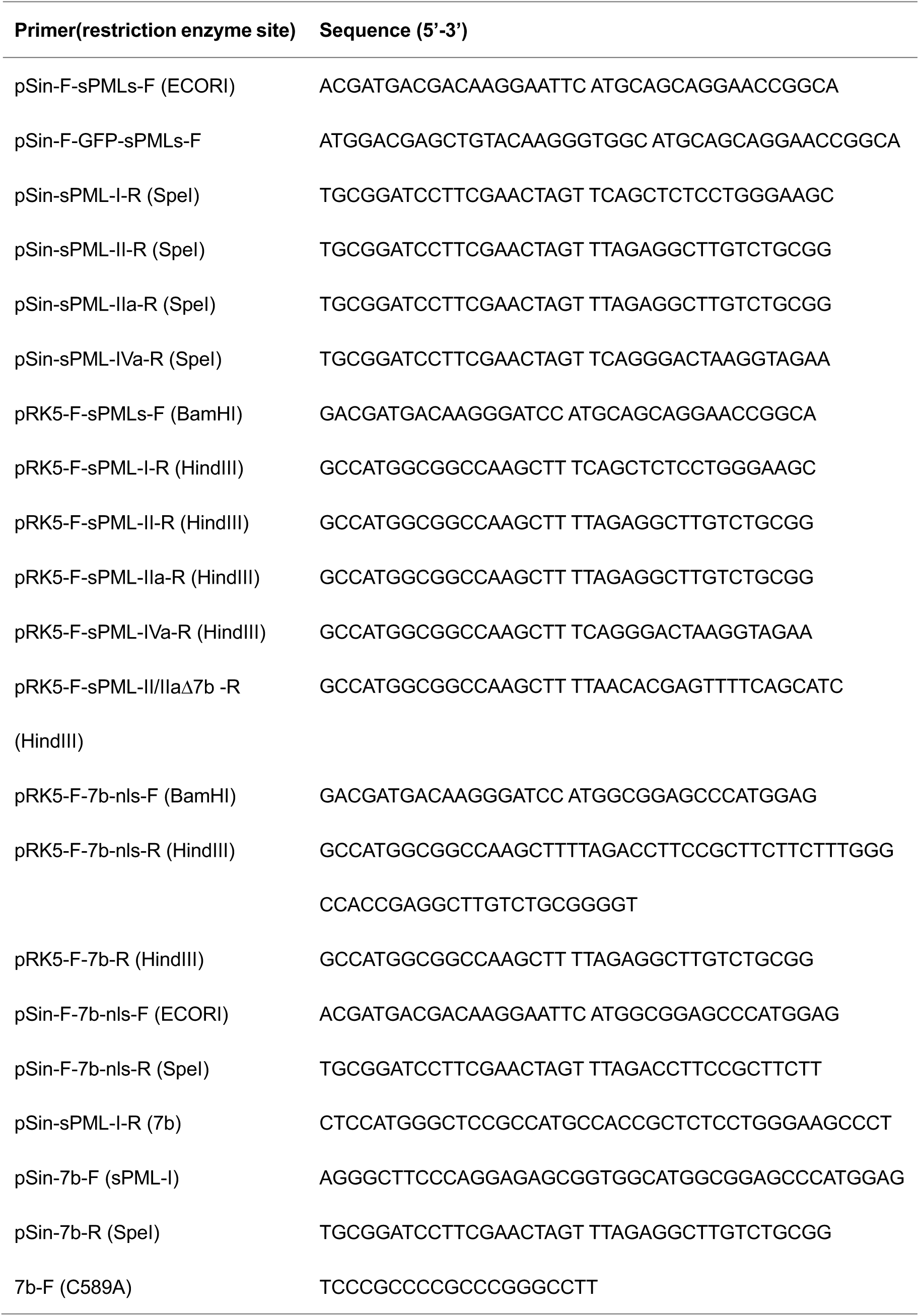
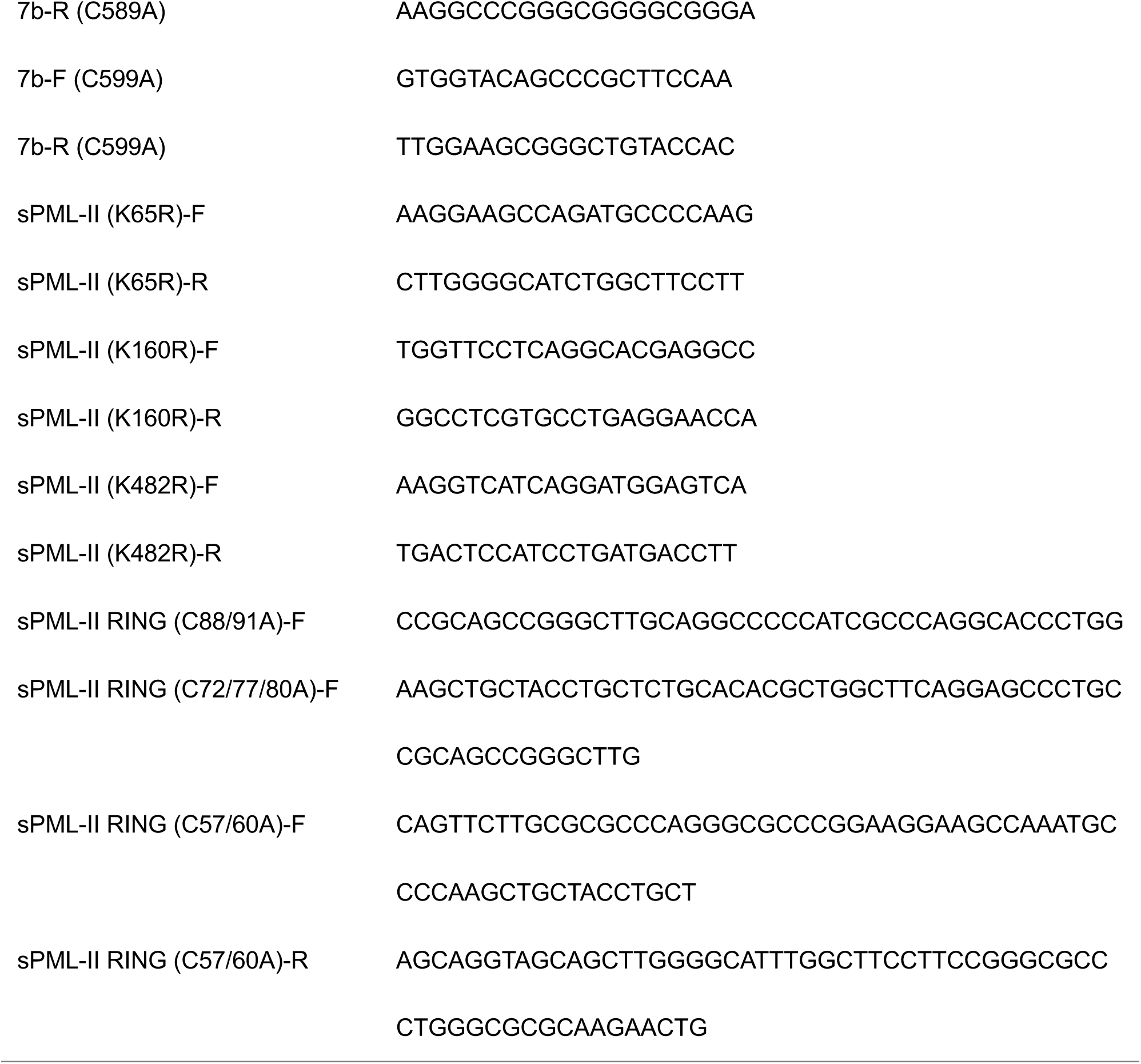
Primers used for sPML isoforms and their mutants cloning

Plasmids were transfected into HEK293T cells using jetPRIME (Polyplus) and PK15 cells using Lipofectamine LTX Reagent (Invitrogen) following the corresponding manufacturer’s protocol.

### Generation of sPML-KO PK15 cells

sPML-KO PK15 cells were generated using TALEN technology using the Fast TALE™ TALEN Assembly Kit (SIDANSAI) according to the manufacturer’s instruction (35). The first exon of PML was targeted by TALEN with the left and right arm sequence of being -GCAGCAGGAACCGGCAC- and - GGGGTCGTCTTGGGGCA-, respectively. PK15 cells were seeded into a 6-well dish and transfected with TALEN plasmids when 70% confluency was achieved. After 24 h of transfection, a medium contained 2.5 μg/ml puromycin was added to the cells. 72 h later, the survived cells were diluted and seeded into a 96-well dish at 0.5 cell/well in a complete DMEM medium. Cells of a single colony were expanded and then examined for sPML expression by immunofluorescent microscopy. sPML-NBs negative cells were further subject to sequencing of exon-1 for verification.

### Generation of stable cell lines

pSin-EF2-puro plasmids encoding aPML-I to -IVa or mutants were transfected into HEK293 cells together with two lentivirus packaging plasmids, pMD2.G, and psPAX2, and the ratio of pSin: psPAX2: pMD2.G was 2:2:1. After 48 h of transfection, the supernatants were harvested, filtered through a 0.45 u syringe filter and then used to infect sPML-WT or sPML-KO PK15 cells in the presence of 8 μg/mL of polybrene. Cells stably expressing sPML isoforms or mutants were selected by treating the infected cells with puromycin at 5 μg/mL for a week.

### Immunofluorescence microscopy

Cells grown on glass coverslips were fixed in 4% paraformaldehyde for 30 min at room temperature and then permeabilized with 0.2% Triton X-100 for 15 min on ice. After washing and blocking in phosphate-buffered saline (PBS) containing 1% bovine serum albumin (BSA) for 30 min, the cells were incubated with specific primary antibodies for 1 h at room temperature followed by FITC or TRITC conjugated secondary antibodies for 30 min. Nuclei were stained with DAPI for 3 to 5 min. Images were captured using a Nikon Eclipse Ni-E microscope or a Leica Wetzlar GmbH microscope. The captured images were processed and analyzed using SPOT software (Nikon).

### Western blot analysis

Whole-cell lysates were prepared in lysis buffer (50 mM Tris-Cl at pH 8.0, 150 mM NaCl, 1.0% Triton X-100, 10% glycerol, 20 mM NaF, 1 mM DTT, and 1× complete protease mixture). The proteins were separated by 10% sodium dodecyl sulfate-polyacrylamide gel electrophoresis (SDS-PAGE) and transferred to nitrocellulose membranes. The membranes were blocked with 5% skim milk in PBST (PBS containing 0.5% Tween 20) for 2 h at room temperature and were then incubated with specific primary antibodies overnight at 4°C followed by secondary antibodies for 45 min at room temperature. The reactive protein bands were visualized using an enhanced chemiluminescence (ECL) reagent with Tanon-5200 luminescent imaging workstation.

### Virus infection and titer determination

PK15 cells were infected with PRV WT or PRV-EP0 KO viruses with the indicated MOI for 1 h followed by washes with PBS and incubation in complete DMEM supplemented with 5% FBS for the indicated durations. The supernatants were collected for titer determination and the remaining cells were used for western blot analysis.

The viral yields of PRV WT or PRV-EP0 KO were determined by plaque assay in Vero cells. Briefly, the collected supernatants from viruses infected PK15 cells were cleared of cell debris by centrifugation, and then used to infect Vero cells in duplicate or triplicate with serial dilutions for 1 h in serum free DMEM. After washes with PBS, the cells were overlaid with 1× DMEM/1% agarose, and incubated at 37 °C until plaque formation was observed (72 h-96 h). The cells were stained with 0.5% neutral red for 4 h-6 h at 37 °C, and the plaques were counted.

### Real-time PCR

Total RNAs were extracted from PK15 cells using TRIzol (Invitrogen) following the manufacturer’s protocol. A total of 0.8 μg RNA from different treatments was reversely transcribed into cDNA using M-MLV reverse transcriptase (Promega) with an oligo (dT) 18 primer. Real-time PCR was performed using an UltraSYBR Mixture (Beijing CoWin Biotech, Beijing, China) on a ViiA 7 real-time PCR system (Applied Biosystems). Viral rRNAs were normalized to swine 28S rRNA expression. Gene-specific primers used for RT–PCR assays included PRV-IE180 forward (5’-ACCACCACCGTCGCCGTCGAGACCGTC-3’) and reverse (5’-GACGGTCTCGACGGCGACGGTGGTGGT-3’), PRV-TK forward (5’-ATGACGGTCGTCTTTGACCGCCAC-3’) and reverse (5’-CGCTGATGTCCCCGACGATGAA-3’), PRV-EP0 forward (5’-GGGTGTGAACTATATCGACACGTC-3’) and reverse (5’-TCAGAGTCAGAGTGTGCCTCG-3’) and swine 28S forward (5’-GGGCCGAAACGATCTCAACC-3’) and reverse (5’-GCCGGGCTTCTTACCCATT-3’) primers.

### Reporter Assay

HEK293T cells were seeded in 24-well plates and transfected with 0.4 μg of 5xGal-TK-luciferase reporter gene plasmid, 50 ng of pCMV-β-galactosidase, and various amounts of plasmids expressing Gal, Gal-7b, Gal-7b(2CA) or Flag-7b. The total amount of DNA was made constant by adding the pRK5 vector. 24 h after transfection, cells were harvested and assayed for luciferase activity with a firefly luciferase system (Promega), according to the manufacturer’s instruction. Luciferase activities were normalized on the basis of the activities of the co-transfected β-galactosidase. Data shown are representative of three independent experiments done in duplicate.

### Statistical analysis

Statistical analyses were performed using GraphPad Prism software to perform Student’s t test or analysis of variance (ANOVA) on at least three independent replicates. P values of <0.05 were considered statistically significant for each test. * *P* < 0.05; ** *P* < 0.01; *** *P* < 0.001.

## RESULTS

### Swine PML-NBs inhibit PRV

To characterize the relationship between PML-NBs and PRV in porcine cells, we first observed sPML-NBs in several types of porcine cells by performing immunofluorescent microscopy using an antibody against sPML. sPML forms NBs as expected in all the cell types examined, and the number of sPML-NBs in freshly isolated primary cells including porcine kidney cells and porcine alveolar macrophages (PAM) was substantially higher than the established cell lines PK15 and CRL (Fig. 1A left and middle panels). This result is consistent with the reported findings that the number of PML-NBs is low in immortalized or certain types of cancer cells due to aberrant signaling in these cells leading to PML destabilization (36, 37). The low number of sPML-NBs in PK15 and CRL cells was dramatically increased by interferon treatment for 12 h (Fig. 1A right panel), indicating that *spml* is also an interferon responsive gene.

**Figure 1.**
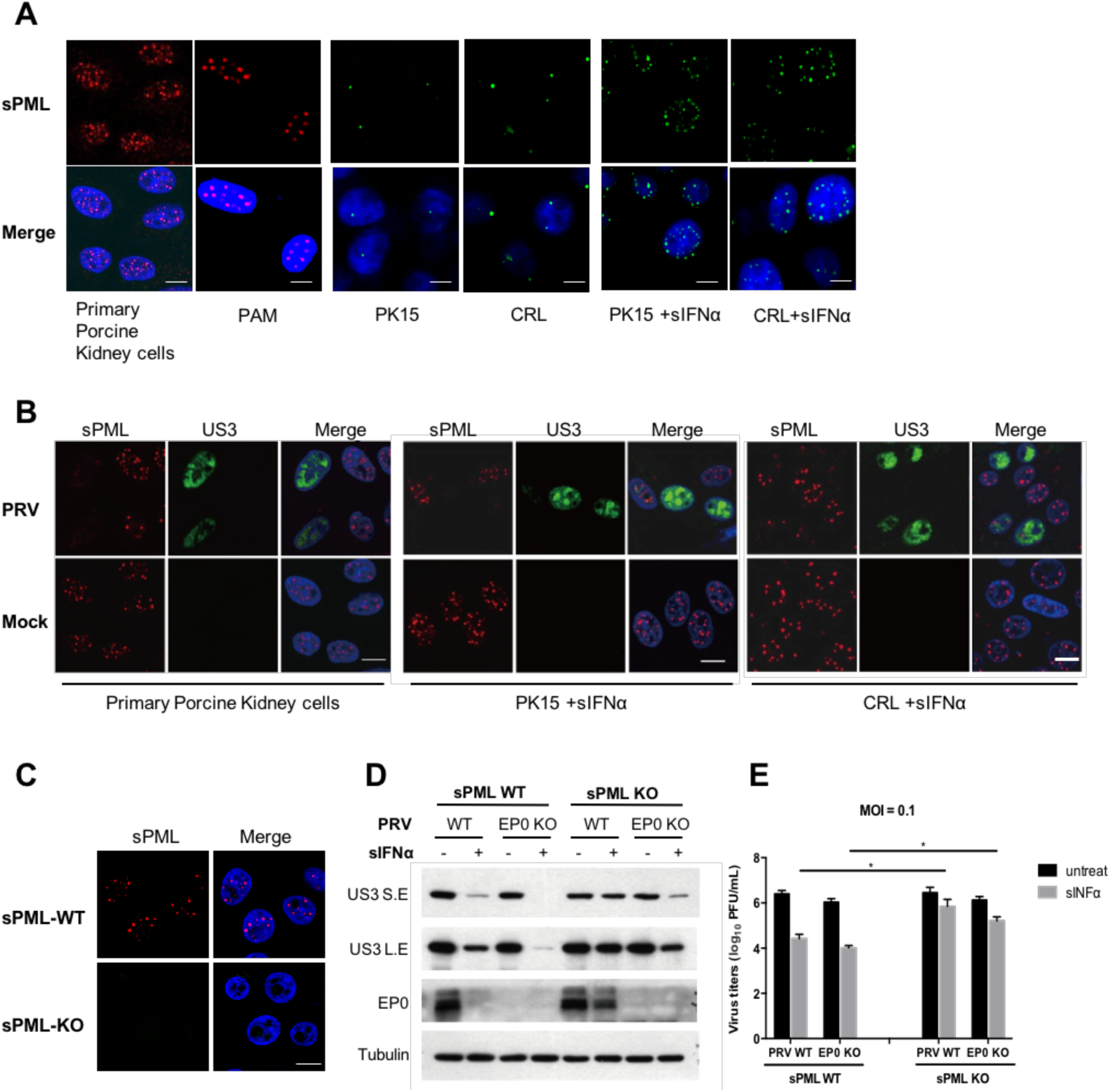
Swine PML-NBs and their relationships with PRV infections. (**A**) Representative immunofluorescent images showing sPML-NBs in several types of porcine cells, including freshly isolated primary porcine kidney cells and porcine alveolar macrophages (PAM) (red, left panel), and cell lines PK15 and CRL without or with 12 h of sIFNα pre-treatment (green, middle and right panels). The cells were stained with an anti-sPML antibody, and the nuclei counterstained with DAPI (blue). Scale bar is 5 μm. (**B**) Representative immunofluorescent images showing the disappearance of sPML-NBs (red) in PRV infected cells (US3 positive, green) as compared to un-infected cells (US3 negative). Primary porcine kidney cells (left), and PK15 (middle) and CRL cells (right) pre-treated with sIFNα were infected with PRV WT (MOI=1) and then stained with a rabbit anti-sPML, and a mouse anti-US3 primary antibody at 24 hpi. The nuclei were stained with DAPI (blue). Scale bar is 5 μm. (**C**) Immunofluorescent microscopy showing that sPML was knocked out in sPML-KO PK15 cells. sPML-WT and -KO PK15 cells were treated with sIFNα (500U/mL) for 12 h before immunostaining with an anti-sPML (red) antibody. The nuclei were stained with DAPI (blue). Scale bar is 5 μm. (**D and E**) Knockout of sPML increases PRV infection in PK15 cells pre-treated with sIFNα. Following treatments with PBS or sIFNα (500U/mL) for 12 h, sPML-WT and KO PK15 cells were infected with PRV-WT or PRV-EP0 KO (MOI=0.1) for 24 h. Cells were collected for western blot analysis of viral protein expressions using the indicated antibodies (D). Viruses released in supernatants were determined by plaque assay (E). Data are shown as mean ± SD of three independent experiments. Statistical analyses were performed by ANOVA, using GraphPad Prism software. *p<0.05.

We then examined the effect of PRV infection on sPML-NBs by infecting cells with PRV for 24 h. PRV infection resulted in disappearance of sPML-NBs in primary porcine kidney cells or PK15 and CRL cells pre-treated with swine IFNa (Fig. 1B), confirming in porcine cells that a-herpesvirus infection disrupts PML-NBs.

Next, we analyzed the effect of knockout of sPML in PK15 cells on PRV infection. sPML was knocked out in PK15 cells by using TALEN technique. Consequently, no sPML fluorescent signal was detected in sPML-knockout (sPML-KO) cells even after interferon treatment (Fig. 1C). sPML-WT and -KO PK15 cells with or without IFN pretreatment were infected with a PRV Bartha wild type (WT) and an EP0 deleted strain (EP0 KO) at MOI 0.1 and then examined viral gene expression by western blotting or infectious viral particle production by measuring the PFU (Fig. 1D and 1E). EP0 is a homolog of ICP0 of HSV-1 known to disrupt PML-NBs (26, 27). When not treated with sIFNa, no significant difference in viral replication for both WT and EP0 KO PRV strains were observed between these two cell lines. These results were not surprising given that the number of sPML-NBs in PK15 cells is remarkably low. Pre-treatment of cells with IFN significantly increased the number of sPML-NBs, and also markedly reduced replications of WT and EP0 KO PRV in PK15 WT cells. IFN also reduced viral replications in sPML-KO cells, but the effects were much less dramatic, indicating that sPML contributed to the anti-viral effect of IFN. Compared with the PRV-WT, EP0 KO viruses were more sensitive to IFN in both cell lines, confirming the previous findings that EP0 is a prominent viral protein that disrupts cellular anti-viral mechanisms such as PML-NBs.

### sPML isoform II and IIa not I and IVa possess an anti-PRV activity

To validate the anti-viral activity of sPML to PRV, we set out to examine the direct effect of expressing sPML on PRV infection by first cloning sPML cDNAs. Analysis of the human and swine *pml* gene indicates *spml* is very similar to human *pml* in terms to exon composition and alternative splicing sites (Fig. 2A), thus, likely generates similar isoforms to those identified in human. Based on the predicated cDNA sequences for the main nuclear sPML isoforms, we successfully cloned four sPML isoforms in this study through PCR using the cDNAs generated from PK15 cells as templates, and designated them as sPML-I, sPML-II, sPML-IIa and sPML-IVa corresponding to each related human PML isoform. sPML-IIa and -IVa are variants of sPML-II and IV, respectively, lacking exon 5 (Fig. 2A). All four sPML isoforms formed typical PML-NBs when expressed in sPML-KO PK15 cells (Fig. 2B).

**Figure 2.**
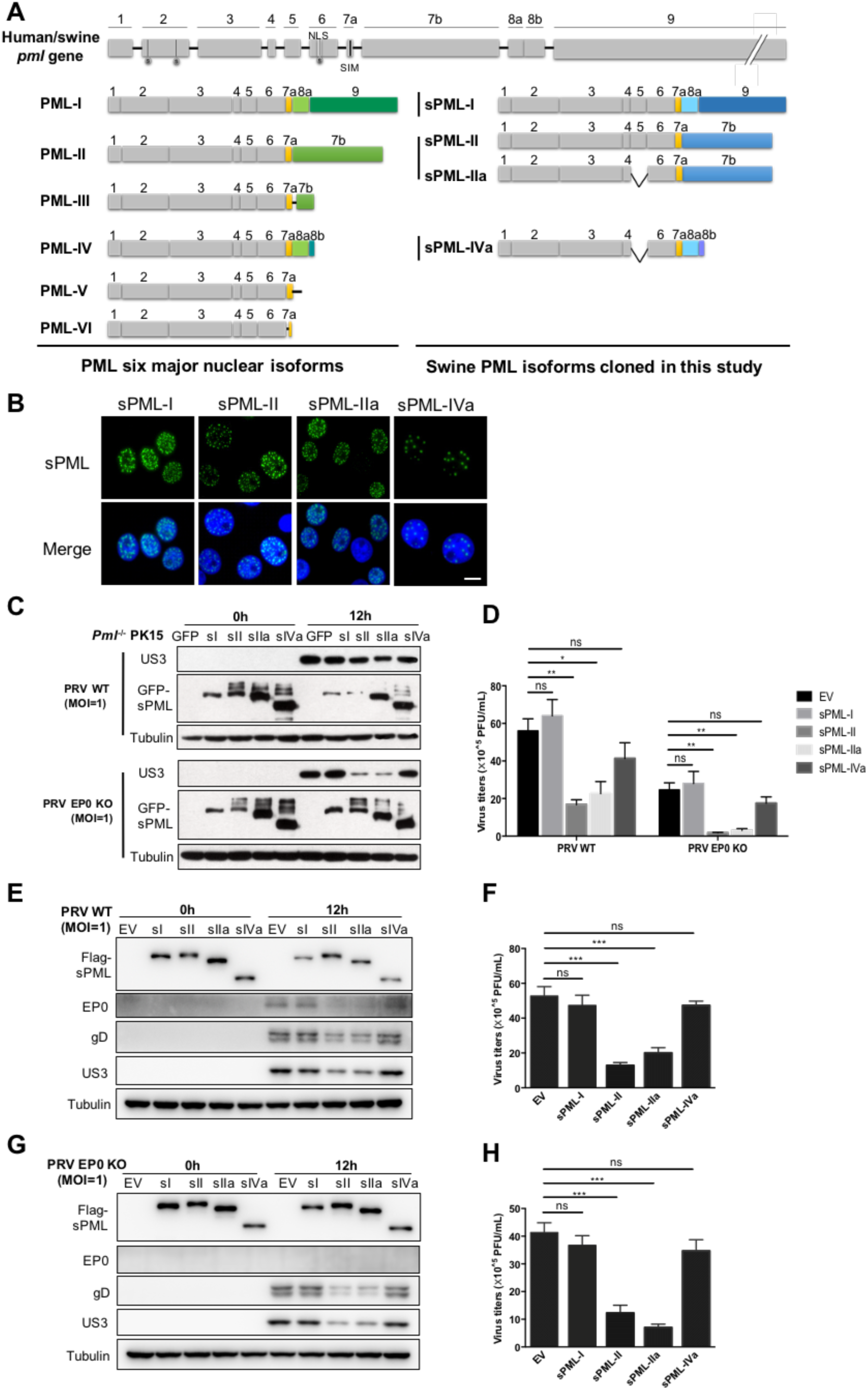
sPML Isoform II/IIa not I and IVa restrict PRV infection. (**A**) Schematic diagrams of human/swine *pml* gene, the six major nuclear isoforms of human PML and likely swine PML (sPML), and the four sPML isoforms cloned in this study. S: SUMO modification, NLS: nuclear location sequence, SIM: SUMO Non-covalent interaction motif. (**B**) Green fluorescent protein (GFP) fluorescence analysis of sPML-NBs in sPML-KO PK15 cells stably expressing GFP-tagged sPML isoforms as indicated. The nuclei were counterstained with DAPI (blue). Scale bar is 5 μm. (**C and D**) sPML-KO PK15 cells stably expressing GFP or GFP-tagged sPML isoforms (sI, sII, sIIa or sIVa) were infected with PRV-WT (MOI=1) or PRV-EP0 KO (MOI=1) for 12 h. Western blotting analyzed US3, GFP-sPML and α-tubulin expression in cell lysates (C). Plaque assay analyzed virus titers in supernatants (D). (**E-H**) sPML-WT PK15 cells transfected with Flag-tagged sPML-I, sPML-II, sPML-IIa, or sPML-IVa expressing plasmids were infected with PRV-WT (MOI=1) (E and F) or PRV-EP0 KO (MOI=1) (G and H) for 12 h. Western blotting analyzed Flag-sPML, EP0, gD, US3 and α-tubulin expression in cell lysates (E and G). Plaque assay analyzed virus titers in supernatants (F and H). Data are shown as mean ± SD of three independent experiments. Statistical analyses were performed by ANOVA, using GraphPad Prism software. *p<0.05; **p<0.01; ***p<0.001.

We then examined whether expressing each sPML isoform in PK15 cells inhibited PRV infection. sPML-KO PK15 cells stably expressing each GFP-sPML isoform as well as GFP were individually established by lentiviral transduction and selection, and the infectivity of these cells to both PRV-WT and -EP0 KO was compared by western analysis of viral protein expressions and viral titer measurements (Fig. 2C and 2D). Compared to GFP expressing cells, sPML-II and -IIa inhibited PRV infections, and to a much more significant degree PRV-EP0 KO infections, whereas sPML-I and -IVa, showed much weaker or even no inhibitions. The similar result was confirmed when F-sPML isoforms were transiently expressed in PK15 cells followed by virus infections (Fig. 2E, 2F, 2G and 2H).

Overall, these data suggest that despite all 4 isoforms forming typical PML-NBs, only sPML-II and –IIa showed strong anti-PRV activity, particularly to PRV-EP0 KO strain. PRV-EP0 KO strain was thus used for most of subsequent studies.

### The unique C-terminal region encoded by exon 7b of sPML-II and -IIa is required for sPML to inhibit PRV infection

As sPML-II and –IIa carry a unique exon 7b region, we asked whether this region possesses an anti-PRV activity. We deleted 7b from F-sPML-II and –IIa (Δ7b), and also constructed a plasmid only expressing 7b region fused with a NLS C-terminally (7b-nls) (Fig. 3A). Adding nls is to ensure 7b is able to enter the nucleus. The anti-PRV activities of these mutants were then analyzed by transfecting the plasmids into PK15 cells followed by PRV-EP0 KO infections. Western blot analysis and viral titer measurements showed that F-sPML-IIΔ7b and –IIaΔ7b completely lost the anti-PRV activity, whereas 7b-nls still strongly inhibited PRV infection, similar to PML-II and -IIa (Fig. 3B and 3C). Additionally, 7b fused to the C-terminal end of GFP-sPML-I (GFP-sPML-I-7b), an isoform without an anti-PRV activity, made the isoform gain the ability to inhibit PRV infection (Fig. 3D, 3E and 3F). This was demonstrated in sPML-KO PK15 cells stably expressing GFP, GFP-sPML-I or GFP-sPML-I-7b. Collectively, these data indicate that 7b possesses an anti-PRV activity, which gives sPML-II/IIa the ability to inhibit PRV infection.

**Figure 3.**
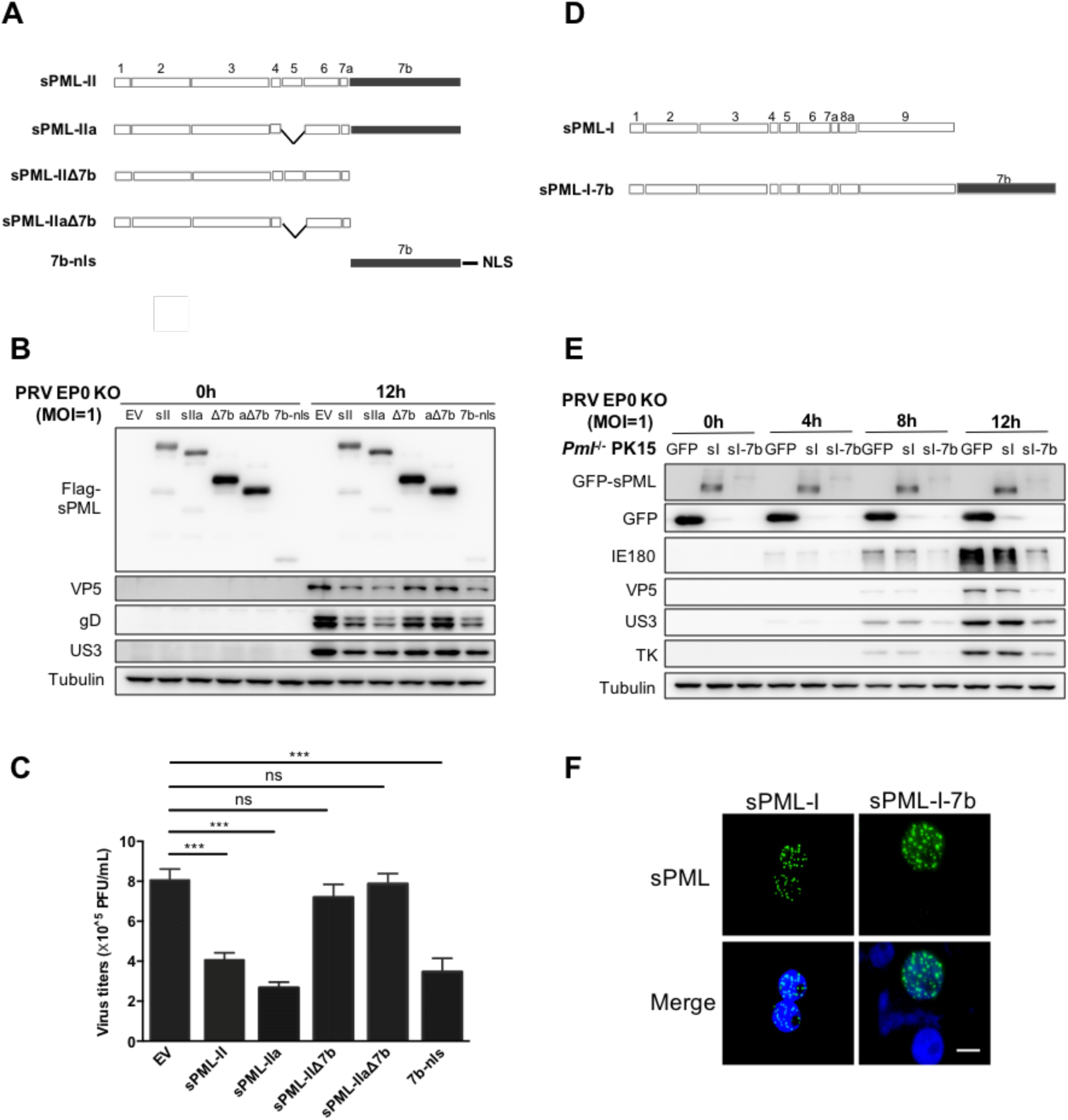
The unique C-terminal region encoded by exon 7b of sPML-II/-IIa possesses an anti-PRV activity. (**A**) Schematic representations of sPML-II/sPML-IIa, 7b deletion mutants sPML-IIΔ7b/sPML-IIaΔ7b, and the nuclear targeting 7b fragment 7b-nls. (**B and C**) 7b region of sPML-II/-IIa possesses an anti-PRV activity. sPML-WT PK15 cells transfected with Flag-sPML-II, -sPML-IIa, -sPML-IIΔ7b (Δ7b), -sPML-IIaΔ7b (aΔ7b) or -7b-nls expressing plasmids were infected with PRV-EP0 KO (MOI=1) for 12 h. Cells were collected for western blot analysis of viral protein expressions using the indicated antibodies (B). Viruses released in supernatants were determined by plaque assay (C). (**D**) Schematic representations of sPML-I and sPML-I-7b in which 7b was fused to the C-terminal end of sPML-I. (**E**) sPML-KO PK15 cells stably expressing the indicated proteins were infected with PRV-EP0 KO (MOI=1) for 4, 8, 12 h. Western blotting analyzed GFP-sPML, GFP, IE180, VP5, US3, TK and α-tubulin expression. (**F**) GFP fluorescence analysis of sPML-NBs in sPML-KO PK15 cells stably expressing GFP-tagged sPML-I or sPML-I-7b. The nuclei were counterstained with DAPI (blue). Scale bar is 5 μm. Data are shown as mean ± SD of three independent experiments. Statistical analyses were performed by ANOVA, using GraphPad Prism software. ***p<0.001.

### Cysteine residue 589 and 599 in 7b is critically involved in the anti-PRV activity of 7b and sPML-II/IIa

To gain more insight into the structural requirement of 7b to inhibit PRV, we analyzed 7b using SMART(38) and revealed that the region of 585-610aa in sPML-II might have a possibility to form a C3H1 type finger despite only two cysteine residues 589 and 599 present (Fig. 4A). To examine whether these two cysteines are involved in the anti-PRV activity of 7b and sPML-II, we mutated them simultaneously or individually into alanine (s) in 7b or sPML-II (Fig. 4A). We then examined the anti-PRV activity of 7b mutants upon transient (Fig. 4B and 4C) and stable expression (Fig. 4D) in PK15 WT cells, and that of sPML-II mutants upon stable expression in sPML-KO PK15 cells (Fig. 4E). Mutation of both cysteines (2CA) or either one (C589A or C599A) completely abolished the anti-PRV activity of 7b or sPML-II (Fig. 4B, 4C, 4D and 4E). These results indicate that cysteine 589 and 599 in 7b is critically involved in the anti-PRV activity of 7b and sPML-II/IIa.

**Figure 4.**
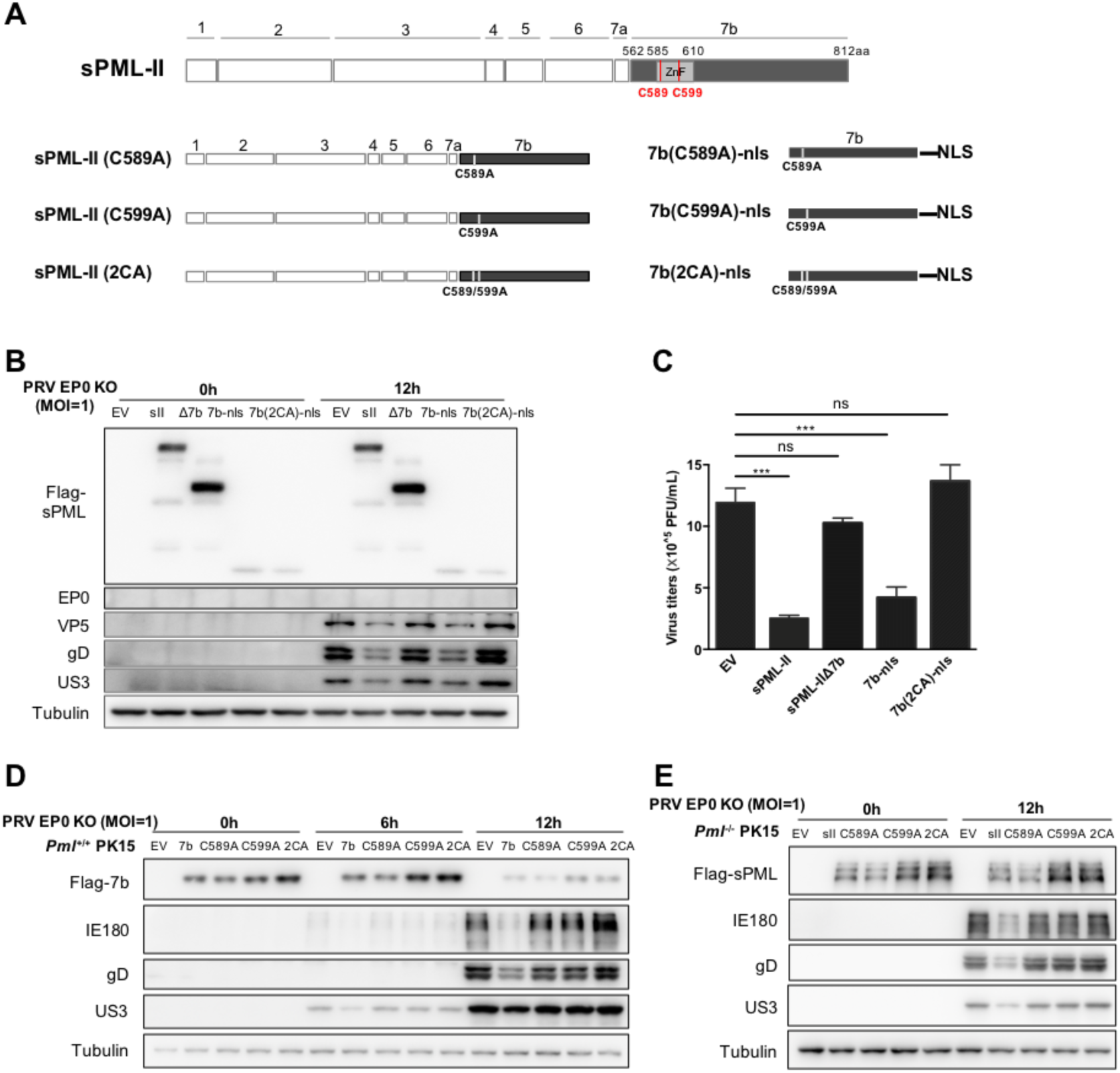
Cysteine residue 589 and 599 in 7b contribute to the anti-PRV activity of 7b and sPML-II/IIa. (**A**) Schematics showing the predicted zinc finger region in 7b using SMART and the mutants used in this study with one or both cysteine residue(s) changed to alanine (**B and C**) sPML-WT PK15 cells expressing sPML-II, or the indicated mutants were infected with PRV-EP0 KO (MOI=1) for 12 h, followed by western blotting (B) and plaque assay (C) as above described. (**D and E**) sPML-WT PK15 cells stably expressing Flag-7b-nls, -7b(C589A)-nls, - 7b(C599A)-nls or -7b(2CA)-nls (D) or sPML-KO PK15 cells stably expressing Flag-sPML-II, -II(C589A), -II(C599A) or -II(2CA) (E) were infected with PRV-EP0 KO (MOI=1) and analyzed by Western blotting. Data are shown as mean ± SD of three independent experiments. Statistical analyses were performed by ANOVA, using GraphPad Prism software. ***p<0.001.

### Localization of 7b in sPML-NB is required for 7b to inhibit PRV infection

The result that expression of just the 7b portion of PML-II in PK15 displays a strong anti-PRV activity is intriguing, raising a question whether formation of sPML-NBs is required for a sPML protein to perform its anti-PRV activity. Since the aforementioned experiments were performed in sPML wild type PK15 cells, we asked whether endogenous sPML-NBs is involved in 7b mediated anti-PRV activity. To test this, we stably expressed F-7b-nls in both sPML-WT and -KO PK15 cells by simultaneously performing lentiviral transductions and selections, and then compared the anti-PRV effect of F-7b-nls in both cell lines, with empty vector and F-7b (2CA)-nls as controls (Fig. 5A, 5B, 5C and 5D). F-7b only exhibited anti-PRV activity in sPML-WT PK15 cells (Fig. 5A and 5B) but no in sPML-KO PK15 (Fig.5C and 5D), suggesting that endogenous sPML-NBs may play a role in facilitating 7b to implement its anti-PRV function. As expected, F-7b(2CA) did not show any inhibition in both cell lines.

**Figure 5.**
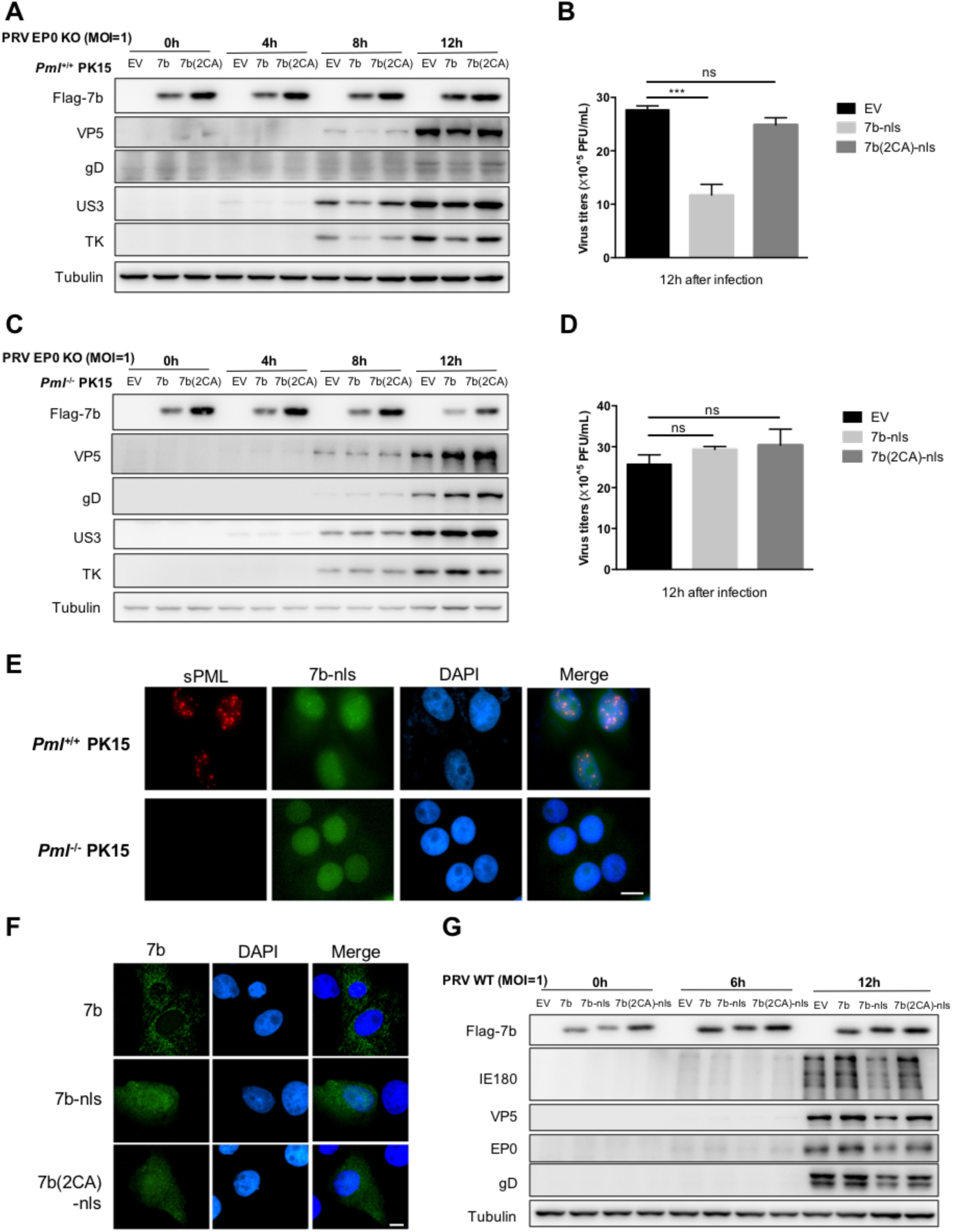
7b mediated inhibition of PRV infection depends on its localization in sPML-NB. (**A-D**) 7b exhibits PRV inhibition in sPML-WT, but not in -KO cells. sPML-WT (A and B) or -KO PK15 (C and D) cells stably expressing Flag-7b-nls or -7b(2CA)-nls were infected with PRV-EP0 KO (MOI=1) for 4, 8 and 12 h, followed by western blot analysis of the cells for viral protein expressions using the indicated antibodies (A and C), and plaque assay of the virus particles released in the supernatants (B and D). (**E**) Representative immunofluorescent images showing the colocalization of 7b (green) and endogenous sPML-NBs (red) in sPML-WT PK15 cells. sPML-WT and -KO PK15 cells stably expressing Flag-tagged 7b-nls were treated with sIFNα (500U/mL) for 12 h before immunostaining with an anti-sPML (red) and anti-Flag (green) antibodies. The nuclei were counterstained with DAPI (blue). Scale bar is 5 μm. (**F**) Representative immunofluorescent images showing that 7b without NLS localizes in the cytoplasm. sPML-WT PK15 cells transfected with Flag-7b, -7b-nls or -7b(2CA)-nls expressing plasmids were analyzed by immunofluorescence microscopy using anti-Flag (green) antibody. The nuclei were counterstained with DAPI (blue). Scale bar is 5 μm. (**G**) 7b without NLS losses the ability to inhibit PRV. sPML-WT PK15 cells transfected with the indicated plasmids were infected with PRV-WT (MOI=1) for 6 and 12 h, followed by western blotting analyzed using the indicated antibodies. Data are shown as mean ± SD of three independent experiments. Statistical analyses were performed by ANOVA, using GraphPad Prism software. ***p<0.001.

To further investigate the connection between sPML-NBs and 7b, we immunostained F-7b in sPML-WT and -KO cells stably expressing F-7b-nls. Confocal microscopy analysis revealed that a portion of F-7b-nls in sPML-WT cells localized in nuclear dot structures, which proved to be sPML-NBs by co-staining with an anti-sPML antibody (Fig. 5E upper panel). In contrast, F-7b-nls in sPML-KO cells was completely diffused in the nucleus (Fig. 5E lower panel). These data suggest that 7b can be recruited into sPML-NBs, and this recruitment may be critical for 7b to execute its anti-PRV function. Supportively, overexpression of F-7b without NLS in sPML-WT PK15 cells, which mainly localized in the cytoplasm, did not show any anti-PRV effect (Fig. 5F and 5G).

Collectively, these data suggest that a functional 7b localizing in sPML-NBs may be required for 7b to perform its anti-PRV function, and endogenous PML-NBs have the ability to recruit it.

### Normal formation of sPML-II NBs is required for sPML-II to inhibit PRV

Next we asked if proper formation of PML-NBs is required for sPML-II to inhibit PRV. PML-NB assembly is initiated by PML N-terminal RBCC region mediated oligomerization, followed by RING domain dependent PML sumoylation, which mainly occurs on sites K65, 160 and 490 (39-41). PML sumoylation critically controls the maturation of PML-NBs (40, 42, 43). We thus examined if abrogation of sPML-II sumoylation affected its anti-PRV activity by either disrupting sPML-II RING or mutating its presumed major sumoylation sites K65, K160 and K482 into arginine. Analysis of the anti-PRV activity of the mutants stably expressed in sPML-KO PK15 cells in comparison with wild type sPML-II showed that the RING inactivation mutant F-sPML-II-RINGca, in which seven cysteine residues of the classic C3HC4 RING finger domain (C57/C60/C72/C77/C80/C88/C91) was mutated into alanine, lost the anti-PRV activity (Fig. 6A). Similarly, sumoylation site mutants sPML-II 2KR (K65/160R) and sPML-II 3KR (K65/160/482R) also lost the anti-PRV activity (Fig. 6C).

**Figure 6.**
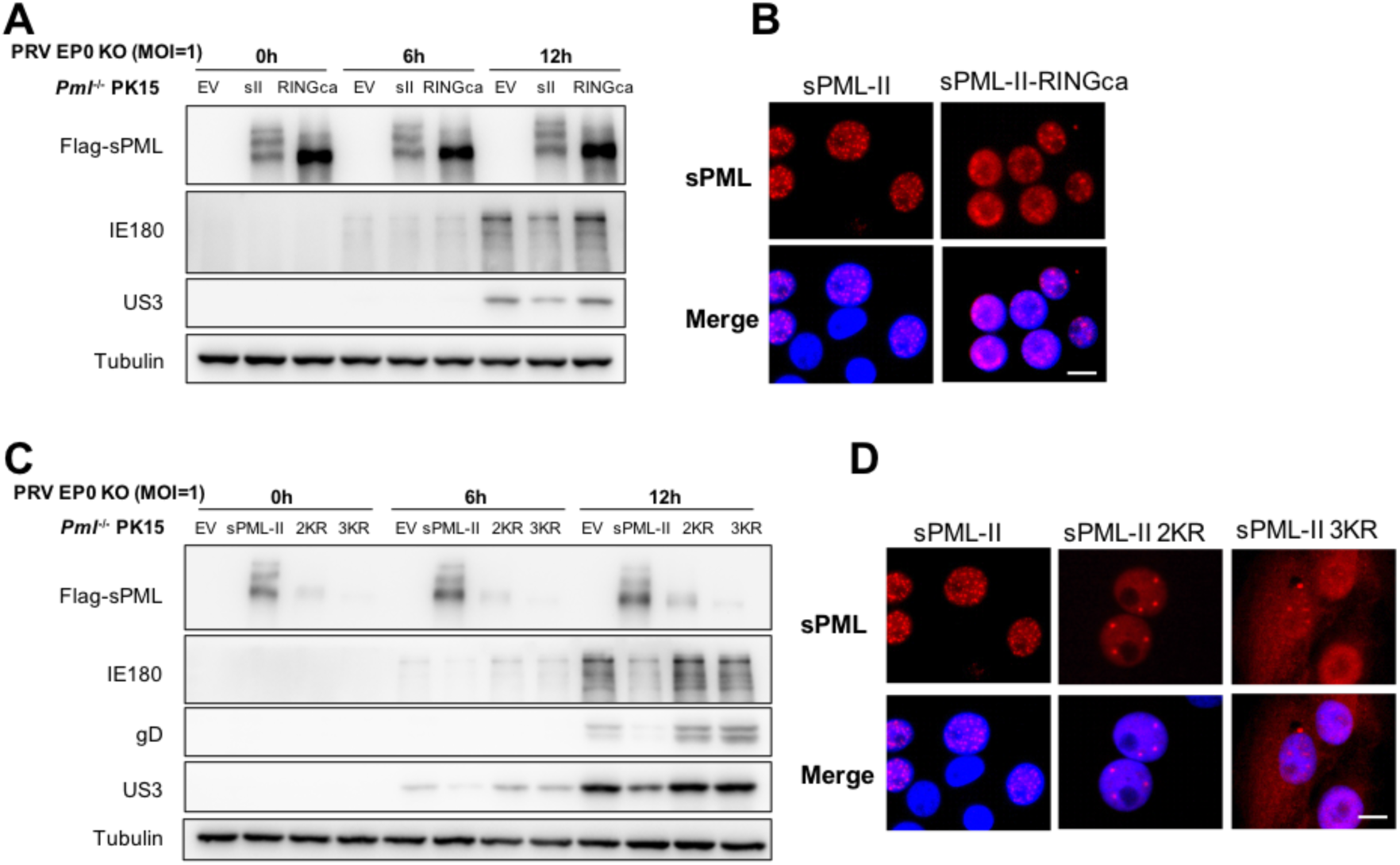
sPML-II inhibits PRV dependently on the normal formation of sPML-NBs. (**A and B**) The RING finger domain is required for sPML-II to inhibit PRV. sPML-KO PK15 cells stably expressing Flag-sPML-II or -sPML-II-RINGca were either infected with PRV-EP0 KO (MOI=1) for 6 and 12 h, followed by western blot analysis using the indicated antibodies (A), or immuno-stained with an anti-Flag (red) antibody followed by immunofluorescence microscopy (B). The nuclei were counterstained with DAPI (blue). Scale bar is 5 μm. (**C and D**) Lysine residues 65, 160 and 482 contribute to the anti-PRV activity of sPML-II. sPML-KO PK15 cells stably expressing Flag-sPML-II, -sPML-II 2KR (K65/160R) or -sPML-II 3KR (K65/160/482R) were either infected with PRV-EP0 KO (MOI=1) for 6 and 12 h, followed by western blot analysis using the indicated antibodies (C), or immuno-stained with an anti-Flag (red) antibody followed by immunofluorescence microscopy (D). The nuclei were counterstained with DAPI (blue). Scale bar is 5 μm.

As expected, the upper modified bands in F-sPML-II expressing cells, which presumably were sumoylated F-sPML-II, disappeared in F-sPML-II-RINGca expressing cells (Fig. 6A), supporting that the RING finger domain critically controls PML sumoylation. However, immunostaining showed that F-sPML-II-RINGca still formed some nuclear dots even though displaying more diffused staining in the nucleoplasm compared with wild type F-sPML-II (Fig. 6B). sPML-II 2KR and 3KR also formed some nuclear dots, but the number was greatly reduced (Fig. 6D). The nuclear dots formed by sPML-II-RINGca, 2KR and 3KR are likely aberrant sPML-NBs (Fig. 6B and 6D). sPML-II 3KR also showed cytoplasmic staining, which is probably due to a defect in nuclear translocation resulted from K482 mutation, which lies in the nuclear localization sequence of sPML 468-482aa, like K490 in human. Intriguingly, sPML-2 2KR and 3KR were difficult to detect by western analysis despite showing normal intensity of immunofluorescent staining (Fig. 6C and 6D).

Altogether, these data show that the normal formation of PML-NBs is required for sPML-II to inhibit PRV infection with PML sumoylation being critically involved. Aberrant sPML-NBs formed by sPML-II due to RING or sumoylation site mutations lost the anti-PRV activity despite the presence of 7b.

### sPML exon 7b inhibits viral gene transcriptions

To explore the mechanism underlying PRV inhibition by sPML-II, we focused on 7b and its role in viral gene transcription because it is the effector region of sPML-II which may possess a putative ring-like structure (44). PK15 cells transfected with EV, F-7b-nls or -7b (2CA)-nls were infected with a high titer of PRV (MOI=5), and the kinetics of PRV viral protein expressions (Fig. 7A) and gene transcriptions (Fig. 7B) were monitored at 2, 4 and 6 hours post-infection (hpi). Western analysis revealed that compared with control cells, viral protein expressions in 7b, but not 7b (2CA), expressing cells were substantially reduced from the time point when detection of these proteins became evident, which was 4 hpi for the immediate early gene IE180 and the early gene (E) US3, and 6 hpi for the late gene gD (Fig. 7A). Viral mRNA measurements indicated relative to the control cells the transcriptions of IE180 and two E genes (EP0 and TK) in 7b expressing cells were also significantly reduced from an earlier time point (2 hpi) when we began to monitor viral mRNA productions (Fig. 7B). These results suggest 7b is clearly involved in transcriptional repression of IE180, and it may also inhibit the transcriptions of other viral genes. However, because the expression of IE180 affects later viral gene transcriptions (25), we cannot rule out the possibility that lower expressions of E and L genes in 7b expressing cells is due to the initial lower IE180 expression in 7b expressing cells. Nevertheless, these results suggest 7b may be involved in inhibition of viral gene transcriptions, particularly IE180.

**Figure 7.**
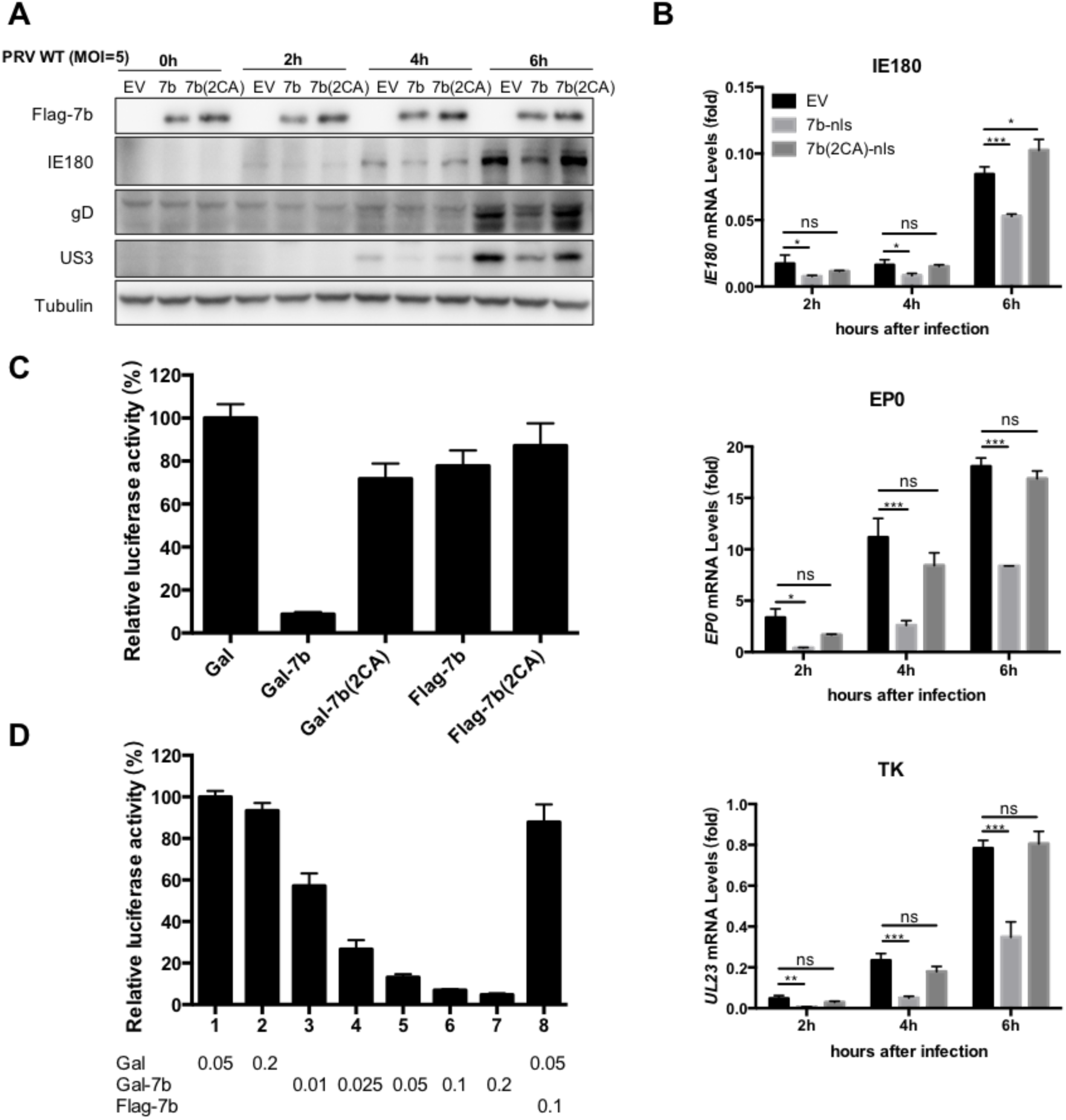
sPML exon 7b function as a transcription repressor. (**A and B**) 7b inhibits viral gene transcriptions during PRV infection. sPML-WT PK15 cells transfected with Flag-7b-nls or -7b(2CA)-nls expressing plasmids were infected with PRV-WT (MOI=5) for 2, 4 and 6 h, followed by western blotting analysis of Flag-7b, IE180, gD, US3 and α-tubulin expression (A), or qRT-PCR analysis of *IE180, EP0* and *UL23* mRNAs (B). (**C and D**) Transcriptional repression by Gal-7b. HEK293T cells were transfected with plasmids expressing Gal, Gal-7b or controls as indicated (C) or with the increased amounts (in micrograms) of Gal or Gal-7b as indicated (D) together with 5xGal-TK-luciferase reporter and CMV-β-galactosidase plasmids. Luciferase activities were normalized with β-galactosidase expression and then compared with that of 50 ng of Gal transfected groups which were arbitrarily set as 100%. Data are shown as mean ± SD of three independent experiments. Statistical analyses were performed by ANOVA, using GraphPad Prism software. *p<0.05; **p<0.01; ***p<0.001.

To direct determine whether 7b possesses a transcriptional repression activity, we fused 7b or 7b (2CA) to the Gal4 DNA-binding domain (Gal-7b or Gal-7b(2CA)) and introduced this fusion into 293T cells together with a luciferase reporter plasmid containing five Gal4 DNA-binding sites upstream of the thymidine kinase (TK) promoter. Gal-7b but not the Gal-7b (2CA) strongly inhibited the luciferase activity (Fig. 7C) and in a dose-dependent manner (Fig. 7D). This transcription repression was dependent on the targeting of 7b to the promoter, as Flag-7b-nls failed to inhibit luciferase activity (Fig. 7C). Thus, exon 7b can function as a transcription repressor when tethered to a promoter.

## DISCUSSION

Here, we demonstrated in swine cells that sPML-NBs inhibit PRV infection in an isoform specific manner. We revealed that only sPML-II/-IIa which carry the unique 7b region can inhibit PRV infection, and that 7b possesses transcriptional repression activity that can suppress viral gene transcriptions at normal sPML-NBs. Our studies not only characterized sPML-NBs in relation to PRV infection for the first time, but also provide a mechanism to explain how sPML-NBs inhibit PRV infection in a sPML-II dependent fashion, which we believe can be extended to explain other scenarios of viral inhibition by a PML isoform.

Our findings suggest that sPML-NBs require at least two properties to inhibit PRV infections: one is the formation of a normal PML-NB with an ability to recruit other molecular, which is RBCC- and sumoylation-dependent; the other is 7b mediated transcriptional repression, which can be provided either as a part of a sPML molecule or by trans. The formation of a functional NB per se is not sufficient but necessary to inhibit PRV infection. sPML molecules with a property to form normal NBs but without a functional 7b element fail to efficiently restrict PRV. These sPML molecules include sPML-I and -IVa, as well sPML-II mutants with either 7b deleted (Δ7b) or transcriptional repression activity abolished (C589A, C599A and 2CA). On the other hand, RING or major sumoylation defective sPML-II mutants also fail to inhibit PRV despite having a functional 7b. Moreover, 7b can only exert its anti-PRV function on the condition of localizing in a functional sPML-NB either as a part of sPML-II, fused with sPML-I or even being recruited there as a separated molecule. These data strongly argue that both normal PML-NBs formation and 7b moiety are required to enable a sPML isoform to restrict PRV infection.

Suppression of viral gene transcription is a critical cellular anti-viral mechanism. It has been reported that numerous proteins are targeted to the HSV-1 viral genome and act coordinately to inhibit viral DNA replication and gene transcription (1, 45, 46). PML-NBs concentrated a number of molecules involved in viral suppression are also recruited to HSV-1 viral DNAs by ATRX or nuclear DNA sensor IFI16 (10, 11, 13). This recruitment process requires a functional RBCC region, PML sumoylation and a SIM motif. In addition to accumulating anti-viral proteins to a viral genome, PML-NBs may also promote various anti-viral processes as a result of multiple dynamic SUMO-SIM interactions, for example Daxx and ATRX meditated epigenetic silencing of viral genomes (2, 5, 47). We speculate that the similar scenario may also exist in swine cells in which sPML-NBs are recruited to PRV genomes allowing the 7b moiety to repress viral gene transcriptions and at the same time promote this process. The observation that 7b can form nuclear dots only in cells with endogenous PML which co-localize with PML-NBs indicates that 7b may also interact with certain component(s) of PML-NBs. Nevertheless, an important contribution of PML-NBs moiety to a-herpesvirus restriction is to recruit gene transcription repressors to viral genomes, and some of the repressors are certain PML isoforms.

sPML-II plays a direct role in repressing PRV gene transcriptions. PML proteins are unique in a sense that splicing variants of all isoforms reportedly co-exist in a NB (36). Increasing evidence indicate that the unique C-terminal moiety of each PML isoform contributes greatly to the formation and diverse functions of PML-NBs. For instance, PML-IV has been extensively studied in the field of cancer biology due to its unique role in binding and regulating the tumor suppressor p53 (48, 49). In the case of HSV-1 infection, only PML-I and -II are reported to partially mediate the anti-HSV-1 functions of PML-NBs, indicating the C-terminal regions of these two isoforms are involved in HSV-1 restriction (24), although the mechanisms by which PML-I and -II suppress HSV-I are not known. We provide evidence to suggest that the mechanism for sPML-II to restrict PRV is to repress viral gene transcriptions mediated by the 7b region. In the presence of 7b, the transcriptions and expressions of all the viral genes examined were suppressed and 7b directly inhibits a reporter gene transcription when tethered to its promoter. Interestingly, homology analysis indicates the c-terminus of PML-I is very conserved between human and swine sharing 74% identity, whereas PML-II 7b is relatively diverse with only 46% identity. Thus, the mechanism by which PML-I and -II restrict HSV-1 might be different from that of sPML-II inhibiting PRV. More comparative studies are required to distinguish the virus specific function of PML isoforms verses the general antiviral property of PML-NBs.

We don’t know the exact mechanism by which 7b inhibits gene transcriptions, but have identified two cysteine residues in a putative zinc finger-like region critically involved in this process. Mutation of either residue resulted in 7b and sPML-II losing the ability to suppress gene transcriptions and/or restrict PRV infection. Although based on the predication the likelihood for the putative zinc finger-like region to form a zinc finger is low, these two cysteine residues certainly play an important role structurally.

## ACKNOWLEDGMENTS

This work was supported by the National Key Research and Development Program of China (grant 2016YFD0500100) and the National Natural Science Foundation of China (grant 31500703).

